# Visuomotor memory is not bound to visual motion

**DOI:** 10.1101/2023.09.27.559701

**Authors:** Matthew Warburton, Carlo Campagnoli, Mark Mon-Williams, Faisal Mushtaq, J. Ryan Morehead

## Abstract

Sensory feedback plays a critical role in motor control and motor learning, with both processes adjusting outgoing motor commands relative to the error between actual sensory feedback and the predictions of a forward model. However, models of motor control rarely specify the exact nature of these predictions. We hypothesized that large differences in low-level perceptual feedback would delineate contextual boundaries for motor memory, as sufficiently different feedback would require a distinct mapping between motor commands and sensory states. We tested this hypothesis by measuring transfer of visuomotor adaptation across contexts where hand movements caused visual motion in opposite directions. Instead of observing that visual feedback is bound to distinct internal models, we found nearly complete transfer of learning across the contexts, with evidence that the motor memory was bound to the planned displacement of the hand, rather than visual features of the task space.

## Introduction

Forward models are thought to play a crucial role in guiding sensorimotor behavior through the prediction of the sensory consequences of movement ^1–3^. These predictions have been shown to be useful for attenuating self-generated sensory signals ^4,5^ and also for robust control of ongoing behavior ^6,7^. In the latter case, feedback control theories propose that behavior is guided by a sensory state estimate that combines observed and predicted sensory feedback ^6,8,9^, with provisions for multiple internal models when the relationship between a motor command and the predicted sensory feedback is context-dependent ^10,11^. However, most models lack clarity on the level at which these sensory predictions operate – for example whether a prediction of movement-contingent visual feedback relies on early visual processing carried out in the primary visual cortex ^12^, complex motion analysis specific to higher visual areas like MT ^13^, or mixed reference frames that combine gaze and body positions ^14^.

This lack of clarity on the characterization of internal model sensory predictions is felt when considering behaviors that differ only in terms of their visual feedback. We recently compared the same center-out reaching task in a mouse *Pointing* context, where a hand movement translates a cursor across a static background, to a first-person shooter (FPS) game inspired *Looking* visual context, where the same movement pans and tilts the view of the environment while the cursor remains central to the screen ^15^.

Movement kinematics and performance in these contexts were highly correlated within individuals, despite profound visual differences, suggesting that both contexts may be making use of the same internal model to plan and correct movements on the fly. If true, this ability for the same internal model to operate across visual feedback contexts cannot be accounted for by prominent models of motor control, which pair forward models of sensory feedback with a movement control policy.

Here we directly probe whether learning is shared between mouse Look and Point feedback by measuring the transfer of visuomotor adaptation between these contexts. Transfer of adaptation is a well-established method to infer properties of internal models ^16–18^. If learning under a visuomotor perturbation, thought to rely on sensory prediction errors ^19,20^, transfers across Point and Look contexts, it will require a reconsideration of current models of sensorimotor control and learning. Specifically, it is unclear how much learning in a forward model ^8,21^ and/or control policy ^22,23^ contributes to sensorimotor adaptation. Critically, adaptation that transfers across mouse Point and Look conditions cannot be explained by either model operating on low-level processing of visual features or on more complex visual motion signals. Such learning would require that internal model adaptation involve a higher, more abstract level of representation such as visuospatial displacement vectors.

To investigate whether internal models are sensitive to the specificity of visual feedback provided for a common movement, we introduced a 30° visuomotor rotation (VMR) between a reaching movement, made with a computer mouse or trackpad, and the visual feedback of the reach, requiring that participants adapt their movements to reinstate good performance ^24,25^ (see supplemental videos). We first compared the adaptive response to the perturbation and the exhibited after-effects, finding nearly identical learning profiles between contexts. Next, we examined transfer of learning between contexts, observing near-complete transfer of learning upon a context switch both when the perturbation was either left on or turned off, confirming the use of a shared internal model across visual contexts. Finally, we found that the learning generalized around the planned movement of the hand, rather than the visual movement of the cursor or the location of the target in the workspace.

## Results

To assess participants’ adaptation to perturbations in an FPS-style experiment, we utilized a framework allowing visual feedback to be compared between and within participants ^15^. Participants used their mouse or trackpad (learning did not differ between input devices across all experiments, so is not considered hereafter) to control a cursor on the screen, appearing like an FPS-style aiming reticle. Upon clicking a start-point, a target immediately appeared, and participants attempted to move to and click on the target (Figure 1a). Computer mouse or trackpad input could control the appearance of the game in two contexts (Figure 1b). In the Point context, movements translated the in-game cursor across a static background. This is consistent with how people normally interact with their computer’s desktop environment, and standard practice for the study of visuomotor adaptation ^24,26–28^. In the Look context, input movements panned and tilted the in-game camera’s view of the virtual environment while the cursor remained central to the screen, consistent with how FPS games are designed. For example, moving the mouse *forward* would cause the in-game character’s view to pan *up*. Critically, the two contexts were equated such that the same input movement in either context would reach a given target, allowing unbiased comparison of behavior between the contexts (see ^15^ for details).

**Figure 1.**
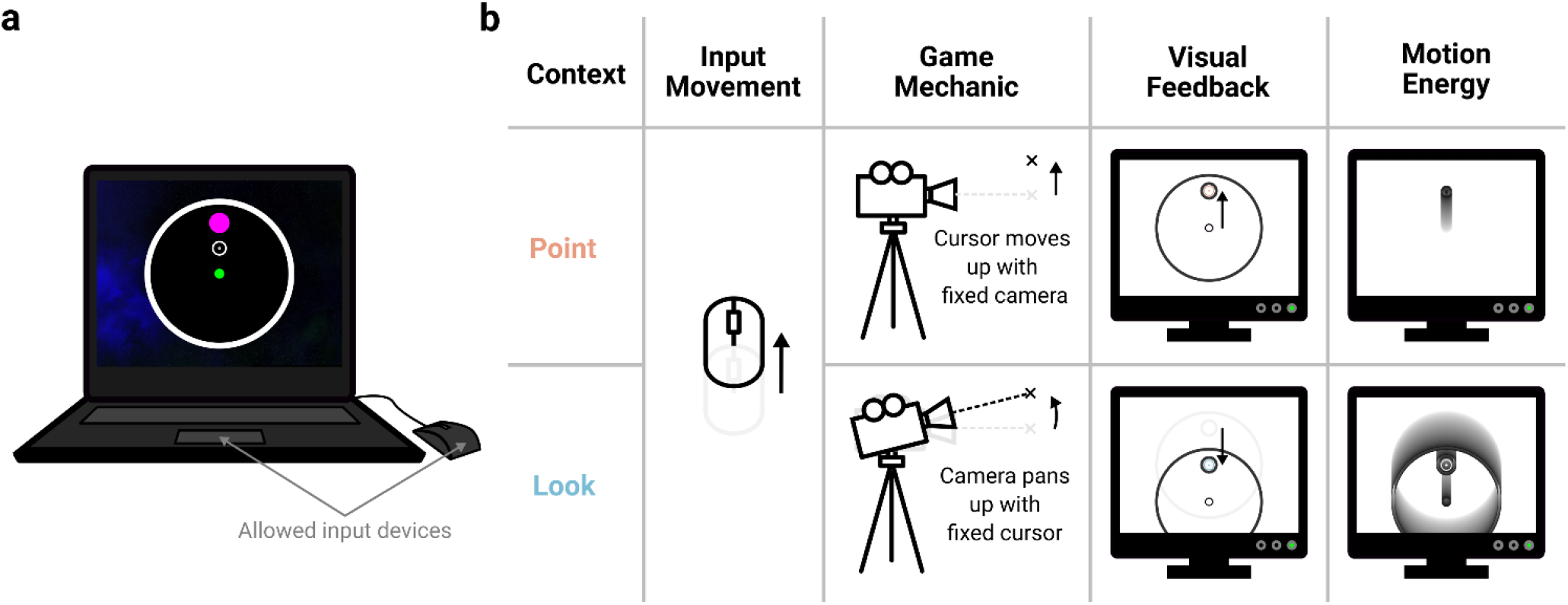
Task setup. **(a)** A visual representation of the task as seen by a participant in the Point mode. Participants used their personal computers and could interact with the task using either a trackpad or computer mouse. Participants left-clicked a start-point and were shown one of four targets to move to. Participants attempted to move to and left-click the target within a continuously staircased time-limit. **(b)** Differences between the contexts, illustrated for a common ‘up’ movement of the input device. In the Point mode, the in-game camera is stationary while a cursor moves across the background, giving visual feedback of only the cursor moving. In the Look mode, the in-game camera pans while the cursor is fixed central to the camera’s view, giving visual feedback where everything except the cursor moves. This gives rise to very different experienced motion energy as the movement evolves over time.

### Participants adapt similarly in the Point and Look contexts

In Experiment 1, we began by assessing whether there were differences in how participants adapted to a perturbation in either context (Figure 2). The perturbation causes different visual errors depending on the context, which the participant must use for both online feedback corrections and trial-to-trial adaptation (Figure 2a). During a trial, upon clicking the start-point, a target immediately appeared in one of four locations, located on an imaginary circle in 90° increments. Participants had to move to and click on the target within a time limit to ‘shoot’ it, otherwise the target would disappear. The time limit was continually staircased throughout the experiment to maintain a ∼50% success rate, inducing the type of time pressure players of FPS games might experience.

**Figure 2.**
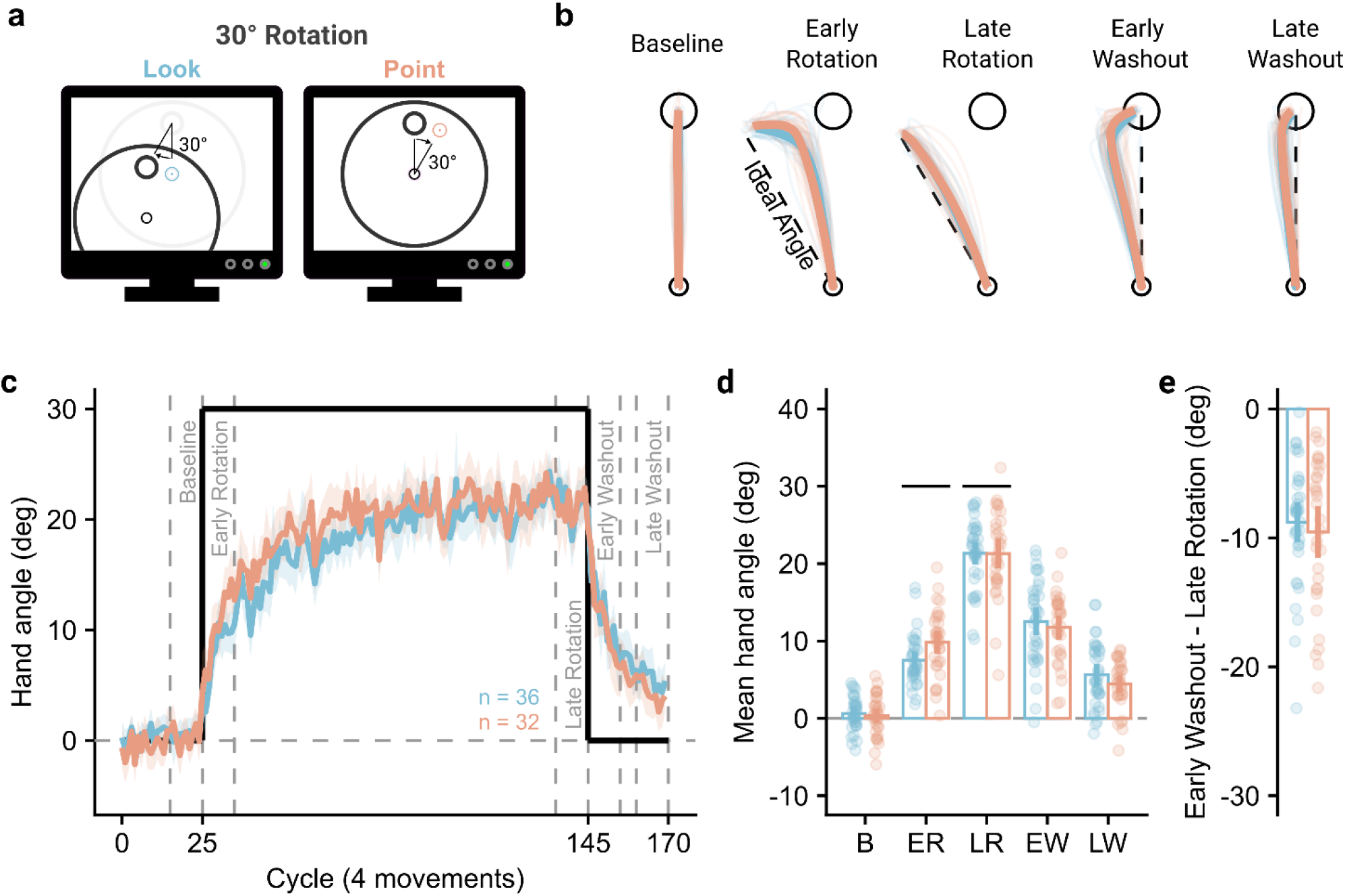
Participants adapted similarly across contexts. **(a)** During the experiment, participants (n = 68) experienced a visuomotor rotation that meant in-game cursor movements were rotated 30° from the veridical movement direction. **(b)** Participant’s hand movements during the perturbation phase reflected compensation for the applied perturbation, with hand paths straightening over the rotation block, and also reflected after-effects, with deviations from baseline reaches at the end of a washout period. Faint lines show individual participants’ average hand path during the period, and thick lines show the group average over participants. **(c)** Time course of the mean hand angle averaged over cycles (4 movements). The line shows the average hand angle from the target direction over participants, with the shaded area showing 95% confidence intervals. The labels show periods of cycles over which measures of learning were operationalised. **(d)** Mean hand angles over periods of the experiment. The period labels are abbreviated from those defined in panel (e). **(e)** Within-participant difference between early washout and asymptotic learning. For panels (f) and (g), points show individual participants, bars show the group mean, and vertical bars show the 95% confidence intervals.

Participants first completed a baseline block with veridical movements, showing average movements that were straight to the target (Figure 2b). A visuomotor perturbation was then introduced, where the visual feedback of a movement was rotated around the start position by 30° from the input movement, requiring participants to compensate for this perturbation to be successful. Early movements (first ten cycles) during this phase showed a large corrective movement near to the target but had straightened by the end of the block (last ten cycles). Once the perturbation ceased, both groups showed similar “hooked” movements in the opposite direction to overcome after-effects, which were still present by the end of the washout period.

This learning is quantified in the average hand angles (measured at peak speed) per group over the time course of the experiment (Figure 2c). During the late rotation period, where learning appeared to have become asymptotic, participants in each group compensated by 21.3° [Point: 95% CI = 19.4° – 23.2°, Look: 19.9° – 22.8°]. To assess differences over the learning block, hand angles over pre-defined periods were assessed using a 2 (context, Point vs Look) x 2 (period, early vs late rotation) mixed ANOVA (Figure 2d).

The ANOVA showed a main effect of period (F(1, 66) = 560.36, p < .001, BF_10_ > 100, η_p_^2^ = .90), indicating participants had greater hand angles during late rotation, but no main effect of context (F(1, 66) = 1.47, p = .230, BF_10_ = 0.51, η_p_^2^ = .02). There was, however, a significant interaction between context and period (F(1, 66) = 4.77, p = .032, BF_10_ = 1.77, η_p_^2^ = .07). Participants in the Point group reached a higher compensatory hand angle than those in the Look group during early learning (estimated marginal mean difference [95% confidence interval] = 2.30° [0.46°, 4.15°], t(66) = 2.49, p = .015, BF_10_ = 3.33), but were not significantly different during late learning (-0.03° [-2.46°, 2.40°], t(66) = -0.03, p = .980 BF_10_ = 0.25). We also characterized the after-effect as the difference between early washout and late learning, to account for any potential differences in asymptotic learning that may have arisen (Figure 2e). The decrease in hand angle from late learning to early washout was not significantly different between the Point and Look groups (-0.74° [-3.29°, 1.81°], t(66) = 0.58, p = .565, BF_10_ = 0.29).

### Learning transfers from a trained context to the other untrained context

Overall, behavior in Experiment 1 was strikingly similar between contexts, reaching similar asymptotic levels and decaying in a similar fashion upon removal of the perturbation. The similarity of learning, as well as the similarity across multiple movement metrics in a previous study ^15^, is consistent with (but insufficient to prove) the use of a shared internal model. To test this more directly, we trained participants to counteract a perturbation in one context before switching to the other, untrained, context. If these contexts do not share an internal model, we would expect poor transfer of learning to the untrained context.

In Experiment 2a, participants were assigned a (trained) context to learn to counteract the perturbation in. An experimental trial mimicked Experiment 1 but followed a different block schedule. Participants performed a pair of unperturbed baseline blocks in both the untrained and trained context before learning to counteract a 30° visuomotor rotation in the trained context. To observe the expression of after-effects following a context switch, participants then performed an unperturbed block in the untrained context, before finally performing an unperturbed block in the trained context.

There appeared to be a greater difference between contexts than in Experiment 1, with greater hand angles for the group training in the Point context (Figure 3a). Hand angles during the learning block (Figure 3b) significantly increased between early and late learning (F(1, 60) = 604.56, p < .001, BF_10_ > 100, η_p_^2^ = .91), and were greater for participants trained in the Point context (F(1, 60) = 13.35, p < .001, BF_10_ = 51.24, η_p_ ^2^ = .18), with no significant interaction (F(1, 60) = 0.32, p = .575, BF_10_ = 0.29, η_p_^2^ = .01).

**Figure 3.**
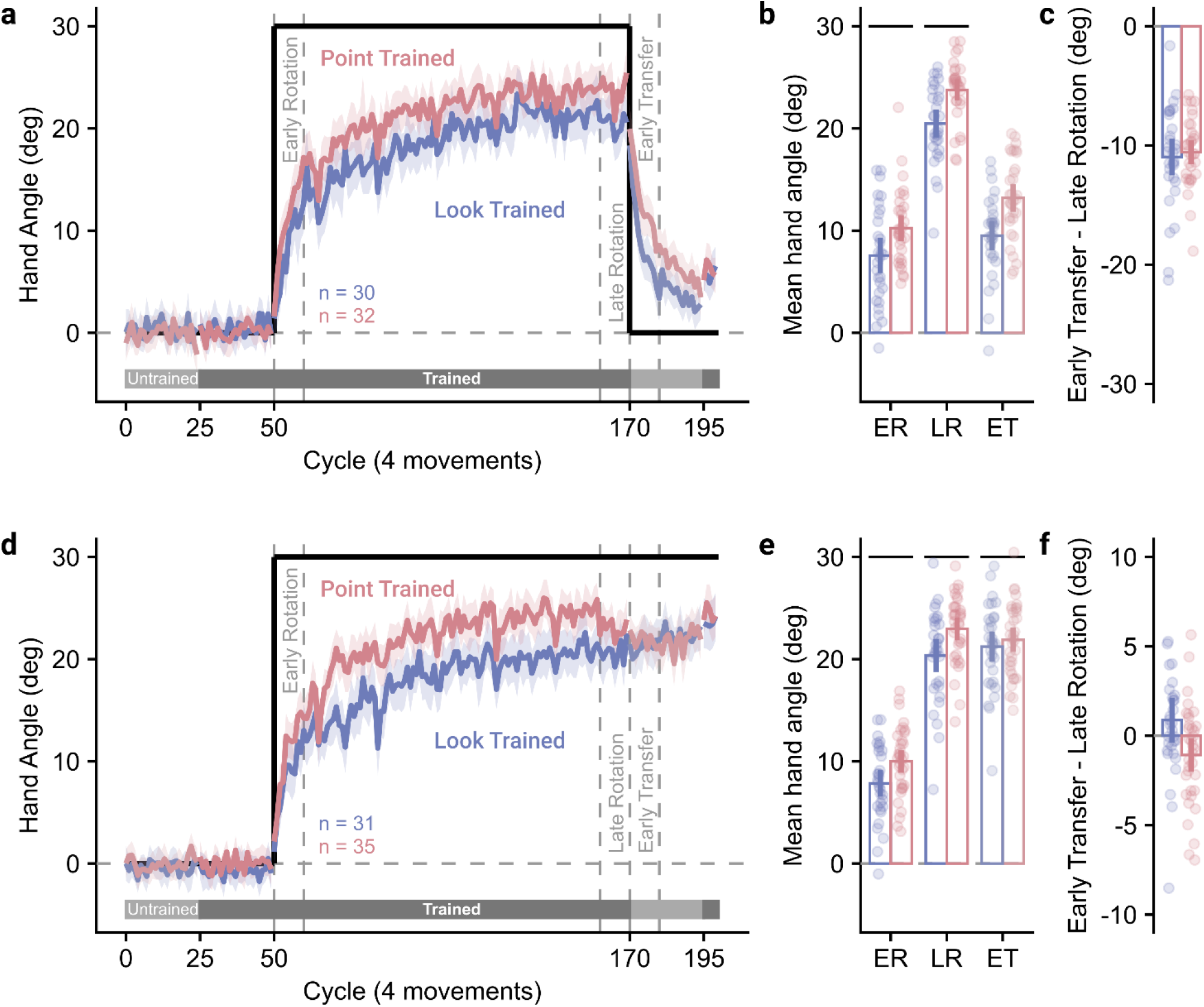
Learning transfers between context. **(a)** Time course of the mean hand angle averaged over cycles (4 movements). The line shows the average hand angle from the target direction over participants, with the shaded area showing 95% confidence intervals. The labels show periods of cycles over which measures of learning were operationalised. Shaded horizontal bars underneath show whether participants were currently completing trials in the trained or untrained context. **(b)** Mean hand angles over periods of the experiment. The period labels are abbreviated from those defined in panel (a). **(c)** Within-participant difference between early transfer and late learning. For panels (b) and (c), points show individual participants, bars show the group mean, and vertical bars show the 95% confidence intervals. **(d-f)** As (a-c) with the perturbation left on when switched to the untrained context.

We next checked for differences in the expression of after-effects. If the learning was bound specifically to the trained context, we would expect a sharp drop in hand angle upon switching to the untrained context. The reduction in hand angle from late learning to early transfer (Figure 3c) was not significantly different between trained contexts (0.41° [-1.45°, 2.27°], t(60) = 0.44, p = .660, BF_10_ = 0.28). Critically, there was no significant difference in the reduction of hand angle when participants switched context, compared to when participants did not switch context in Experiment 1, for either the group trained in the Point context (-1.02° [-3.31°, 1.28°], t(62) = 0.88, p = .380, BF_10_ = 0.36) or Look context (-2.16° [-4.40°, 0.07°], t(64) = 1.93, p = .058, BF_10_ = 1.21). This indicates there was no evidence for a reduction in after-effects when participants switched to an untrained context.

Similar analyses were performed for Experiment 2b, where the perturbation was left on as participants were switched to the untrained context. Hand angles again appeared to be greater for those trained in the Point context (Figure 3d). Hand angles during the learning block (Figure 3e) were again significantly greater during late learning (F(1, 64) = 557.21, p < .001, BF_10_ > 100, η_p_ ^2^ = .90), and for the participants trained in the Point context (F(1, 64) = 10.02, p = .002, BF_10_ = 14.01, η_p_ ^2^ = .14), with no significant interaction (F(1, 64) = 0.18, p = .674, BF_10_ = 0.27, η_p_ ^2^ < .01).

Despite the perturbation being left on in this task, we would still expect a substantial drop in hand angle immediately after transfer if the contexts did not share an internal model. The change in hand angle from late learning to early transfer (Figure 3f) was significantly lower for participants who trained in the Point context compared to the Look context (-1.95 [-3.53°, -0.37°], t(64) = -2.47, p = .016, BF_10_ = 3.18). Participants who trained in the Point context showed a small but significant reduction in hand angle from late learning to early transfer (-1.07 [-2.05°, -0.09°], t(34) = -2.23, p = .032, BF_10_ = 1.56), whereas participants who trained in the Look context showed no significant difference in hand angle between late learning and early transfer (0.88 [-0.43°, 2.19°], t(30) = 1.37, p = .180, BF_10_ = 0.45).

Mixed-effect linear regressions across the untrained transfer block (cycles 170-195) showed that participants trained in the Point context had an intercept significantly lower than asymptotic learning upon switching to the Look context (β = -1.28 [-2.33°, -0.24°], t = -2.41, p = .019), with no significant change over cycles (β = 0.01 [-0.03°, 0.05°], t = 0.53, p = .599), whereas the intercept did not differ for participants trained in the Look context (β = 0.11 [-0.82°, 1.05°], t = 0.23, p = .816) but hand angle did significantly increased over cycles (β = 0.07 [0.03°, 0.12°], t = 3.15, p = .002). Both regressions suggest asymptotic learning is slightly lower in the Look context – with those transferred to it experiencing a rapid decrease in hand angle and those transferred from it able to increase their compensatory angle slightly. Nevertheless, both experiments clearly show that the bulk of learning transfers to the untrained context.

### Learning generalizes around the planned movement of the hand

The previous experiment found that the bulk of learning transfers between the visual contexts. We suggest this shows the contexts do share a common internal model, and that learning generalizes around a feature common to both tasks. Visuomotor rotations induce learning that generalizes locally around an aiming vector ^29,30^. Whether learning is bound to an aim location in visual or motor space is unclear – the two are typically confounded. To investigate this, we used a manipulation taken from the FPS gaming domain, where a subset of games and gamers ‘invert’ their mapping, such that they move their mouse or joystick *forward* to look *down* in the game (Figure 4a). The Inverted Look mapping should therefore allow us to dissociate whether learning at a single target (Figure 4b) generalizes locally around the planned vector of the hand, where it would generalize roughly around the trained handspace direction; or the planned vector of the cursor, where it would generalize roughly around the trained visual target (Figure 4c).

**Figure 4.**
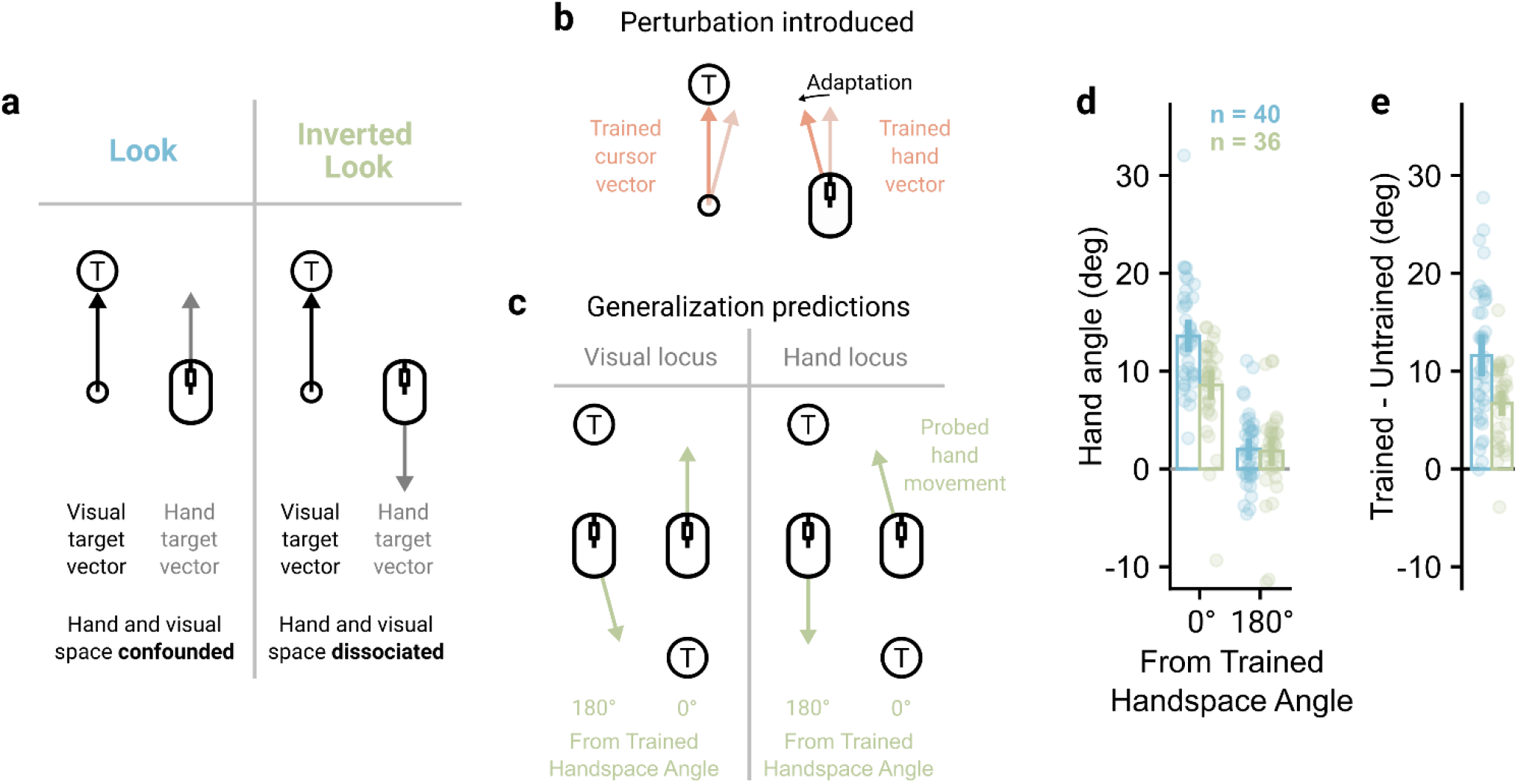
Learning generalizes around the planned hand vector. **(a)** Experiment 3 included an Inverted Look context. When participants move their mouse forward, their in-game view pans down. **(b)** Participants learned to counteract a visuomotor rotation at a single target in the Point context, where the planned cursor and hand vectors were common. **(c)** The inverted look allows the planned cursor and hand vectors to be dissociated. A visual locus of learning should show adaptation when the planned cursor vector is congruent with that trained, whereas a hand-based locus should show adaptation when the planned hand vector is congruent with that trained. **(d)** Participants in both groups showed larger amounts of generalization to the trained movement of the hand. **(e)** Within-participant comparisons between trained and untrained movement direction showed larger hand angles for the trained movement. For panels (d-e), bars show mean hand angles and vertical lines show 95% confidence intervals.

In Experiment 3, participants were split into two groups where the spatial generalization of learning was assessed with either Look or Inverted Look probes. Experimental trials using the probed context proceeded as in the previous experiments with two exceptions – there was no time limit, and the upcoming context was signaled by a color cue as participants returned to the start-point. Experimental trials using the Point context were similar except no feedback was provided as participants moved their hand back to the start-point (see methods). Following baseline trials to introduce this change in trial feedback, participants completed a baseline generalization block, where 15 generalization cycles were performed. Each cycle had three trials in the probed context, first to either the left or right target and then to both the top and bottom targets, followed by 15 Point trials to the top target. No perturbation was applied during this block, and it allowed the later generalization block to be baseline-corrected, removing systematic biases in reaches ^31^.

Participants next experienced 120 Point trials to the top target, where they learned to counteract a 30° visuomotor rotation. Critically, because participants did not have return feedback during these trials, learning should only generalize locally around the aiming vector. Both groups learned to counteract the perturbation similarly, with hand angles increasing between early and late learning (F(1, 74) = 251.69, p < .001, BF_10_ > 100, η_p_ ^2^ = .77), and no significant effect of probed context (F(1, 74) = 0.96, p = .331, BF_10_ = 0.35, η_p_ ^2^ = .01) or interaction (F(1, 74) = 0.04, p = .849, BF_10_ = 0.24, η_p_ ^2^ < .01).

Participants then completed the generalization block, which was identical to the baseline generalization block except that the 15 Point trials per cycle had the 30° visuomotor rotation applied. Because unlearning will occur during the unperturbed probe trials, enough relearning trials (established in a pilot study) were allowed to ‘top up’ learning for the top target. Taking just the probe trials in the same and opposite direction to the trained handspace direction (Figure 4d), a significant main effect of angle from the trained handspace direction indicated learning generalized around the planned hand vector (F(1, 74) = 197.04, p < .001, BF_10_ > 100, η_p_^2^ = .73). Further, there was a main effect of probed context (F(1, 74) = 9.64, p = .003, BF_10_ = 9.66, η_p_ ^2^ = .12), indicating hand angle was greater for the Look probes, and a significant interaction (F(1, 74) = 13.88, p < .001, BF_10_ = 95.26, η_p_ ^2^ = .16). This interaction was followed up by assessing the differences in hand angle between trained and untrained movement directions. Hand angle was greater for the trained handspace direction compared to the untrained handspace direction in both Look (11.61° [9.81°, 13.40°], t(74) = 12.90, p < .001, BF_10_ > 100) and Inverted Look groups (6.74° [4.85°, 8.63°], t(74) = 7.11, p < .001, BF_10_ > 100; Figure 4e). Therefore, both groups show a robust transfer of learning from the trained Point context to the probed Look or Inverted Look context, supporting the previous experiment’s results, and the results are consistent in that learning generalizes around planned movement of the hand.

This difference was, however, significantly greater for the Look group compared to the Inverted Look group (4.87° [2.26°, 7.47°], t(74) = 3.73, p < .001, BF_10_ = 69.94). A mixed-effect linear regression, assessing how hand angle for the trained input direction changed over generalization cycle, showed there was no significant difference between probed context on the first generalization cycle (β = -2.17 [-5.02°, 0.69°], t = -1.48, p = .140). The Look group showed no significant change in hand angle over cycles (β = 0.04 [-0.12°, 0.20°], t = 0.45, p = .651), but the Inverted Look had a significant decrease in hand angle over cycles (β = - 0.29 [-0.53°, -0.05°], t = -2.40, p = .017). Therefore, while hand angle is not significantly lower for the Inverted Look group at the start of the generalization block, the groups diverged over the cycles. This could indicate that, for the Inverted Look group, transferred learning itself reduced over cycles, which could happen if the number of top-up trials was not enough for the group to re-establish a consistent amount of implicit learning, or that participants formed an explicit strategy to overcome the implicit learning shown in the trained input direction, which was possible given participants had full vision during the probe trials.

## Discussion

Here we examined transfer of visuomotor adaptation across two task contexts with movement-contingent visual feedback that was categorically distinct, but with matched motoric elements. We demonstrated that adaptation proceeds similarly in these two contexts, and that this learning readily transfers between the contexts. Finally, we showed that learning generalizes around the planned movement of the hand, rather than any visual feature of the task. These results demonstrate that visuomotor adaptation modifies an element of the visuomotor control circuit that is downstream of the specific visual features of the reaching task. We believe these results have meaningful implications for the theory of internal models for visuomotor control, specifically regarding the specificity of sensory feedback in planning and error correction. Our results are incompatible with models that employ low-level visual information or even complex motion information as an input or a cue for memory retrieval. Instead, our data suggests that the critical factor binding sensory error to motor memory is the planned displacement of the hand in space. The same planned displacement of the hand operating across these visual contexts would also suggest the shared use of internal models operating on higher, abstracted representations of sensory feedback.

### Visual contingencies

By utilizing two visual feedback contexts that deliver motion energy in opposite directions, we found that visuomotor adaptation is not critically tied to the specific details of movement-related visual feedback. Learning progressed similarly between visual contexts in the first experiment, with both groups showing a comparable reduction in hand angle once the visuomotor rotation ceased. In Experiment 2, we showed that learning in one visual context transfers almost entirely to the other, untrained, visual context – when the perturbation was turned off upon a context switch, participants showed a similar decay profile to Experiment 1; and when the perturbation was left on upon a context switch, participants showed no substantial change in hand angle. The third experiment further demonstrated transfer of learning from the trained Point context to untrained Look and Inverted Look contexts. While there is debate around the error signal important for visuomotor adaptation ^20,32,33^, and what changes this error drives ^8,22^, we find that the same motor memories are engaged and altered by these very different forms of visual feedback.

### Sensory predictions

Internal models are ubiquitous in modern theories of motor control and learning ^1–3^. The sensory predictions generated by forward models can address a number of issues ^6^, for example distinguishing self and externally-generated sensory input ^34^, stabilizing state estimates from noisy sensory input ^35^, overcoming the long time-delays associated with sensory processing for real-time feedback corrections ^36^, and guiding learning ^37^. In some cases the details around the sensory predictions are exceptionally well-defined, for example in work assessing the cancellation of self-generated sensory feedback in the weakly electric fish ^5^. In general, however, the output of the forward model is typically stated to be a prediction of the sensory consequences of a movement, without specifying what form that may take.

Based on our current and previous results ^15^, it seems unlikely that sensory predictions are at the level of low-level visual features. Under such a scenario, both visual contexts would require distinct internal models. While models based on a mixture-of-experts ^10,11^ could accommodate this, by distributing learning from one context across others, it would require that participants already had the requisite internal models for the Look context. Only 19 of 68 participants in Experiment 1 reported playing FPS games, with 14 report playing no games at all, and tasks thought to elicit *de novo* acquisition of a new internal model typically show extended periods of poor performance ^38^. It, therefore, seems unlikely participants with little to no experience with FPS games could acquire such internal models and switch between them so rapidly.

Instead, we suggest that sensory predictions are abstracted from the exact visual feedback expected. While work has emphasized the role of the forward model is to predict the *state* of the body and world ^39^, this is unsatisfying as state variables for the cursor like its position and velocity never change in *extrinsic* space for the Look context, as it is clamped to the screen center, so only vary relative to other features in the scene. Further, unlike traditional tablet-based setups, there is no one-to-one mapping between hand and cursor position in the Point context when moving an optical mouse or trackpad – only the movement vector between initial and final hand and cursor positions correlate.

Previous work has found that movement vectors play an important role in motor control and planning. Invariant features of goal-directed reaches, like their straightness in extrinsic space and single-peaked tangential velocity, suggest movements are planned in terms of task-space trajectories ^40–43^. The direction and extent of the trajectory, identifying a movement vector in polar coordinates, appear to be relevant parameters that are specified separately during planning ^44,45^, centered on the effector’s starting position ^46,47^. During adaptation to a visuomotor rotation, the movement vector is the predominant feature remapped ^25,48,49^, with spatial generalization greatest in the direction participants aimed towards ^29,30,50^. By dissociating hand and cursor movements in Experiment 3, we showed that the locus of learning is specifically the planned movement vector of the hand, rather than a visually referenced aim location. The transfer of learning between the two visual contexts in Experiments 2 and 3 would also suggest that the sensory predictions of the consequences of such movements are represented at the level of movement vectors, allowing common internal models to be engaged.

### Contextual cues

Recent work has found effective contextual cues that allow multiple mappings between movements and sensory feedback to be learned concurrently. Participants can learn to counteract opposite signed perturbations, which would normally interfere ^51,52^, when paired with contextual cues relevant to the movement, for example distinct physical workspace locations ^53,54^, planned movements ^55^, or (planned) lead-in or follow-through movements ^52,56–58^. Cues related to visual aspects of the movement, like the visual workspace location that feedback is displayed in ^53^ or where participant gaze is directed ^59^, also appear effective. The visual contexts studied here may, therefore, serve as effective contextual cues, given visual feedback of the movement progresses differently over time and maintaining fixation on the target would require different gaze patterns between the contexts. In Experiments 2 and 3, we did not train participants to associate different visual contexts with different perturbations concurrently, instead finding that, without a reason to differentiate learning, blocked learning in one context showed near-complete transfer to the other.

The difficulty with using the visual contexts as presented here as effective contextual cues is that, prior to movement onset, the two appear identical. This means additional arbitrary cues would have to be used, like the ring color used for Experiment 3, but such cues are generally ineffective at allowing opposing perturbations to be learned ^51,53,60^. However, we do know that a subset of gamers express an Inverted Look mapping in the context of an FPS game, moving their mouse forward to reach a target directly down, yet switch to typical Point movements when moving their cursor down on the in-game menu or in other non-FPS games. Future research could look at the time scale and cues necessary to allow opposing mappings to be retained long-term and contextually switched in this manner.

### Studying learning in FPS games

Video games offer an exciting opportunity to study situated skilled behavior by simply monitoring the inputs, typically from the mouse and keyboard, that are used to interact with the game, allowing relatively passive large-scale data collection. For example, data collected from a commercial FPS aim-trainer game, which uses tasks isolating specific behaviors like aiming, has been used to investigate the long-term effects of practice ^61^ and the kinematic correlates of skill ^62^, and actual gameplay data has been analyzed to understand behavioral differences between novice and professional players ^63^. Such approaches can allow the development a player’s game skill to be tracked ^64,65^, offering insights about long-term motor learning that are hard or impossible to make with restricted laboratory studies. Movement kinematics also inform us about more than just the motor system – they have been used to study the dynamics of a wide range of perceptual and cognitive processes ^66–69^. Given that FPS games typically engage a range of perceptual, cognitive, and motor abilities ^70^, the kinematic data recorded in actual gameplay may therefore provide insights across a number of domains. Here we utilized the differences in how visual feedback is presented in FPS games compared to more traditional pointing style feedback, finding that visuomotor adaptation proceeded similarly and transferred across visual contexts. This, combined with nearly identical kinematic features between the two visual contexts ^15^, should give researchers the confidence to utilize this exciting medium to study skilled behavior.

## Methods

### Participants

In total, 276 participants were recruited to take part in the experiments. After exclusions (n = 2 in Experiment 2: 1 participant recorded no successful movements after the first block, 1 participant’s data did not upload; n = 2 in Experiment 3: participant data did not upload), the final sample was 272 participants. Participants were recruited through Prolific.co, an online recruitment platform, and were restricted to those living in the UK or USA, who had English as a first language, and had a Prolific approval rating of 95% or above. Participants were paid between £6.25 - £8 upon completion for expected completion times between 50 – 60 minutes depending on experiment. Experiments were approved by the School of Psychology Ethics Committee at the University of Leeds, and participants gave informed consent via a web form prior to starting the study. Given the experiment was completed online, which has been associated with noisier responses within and between participants on similar paradigms ^71^, we recruited more participants per group (between 30 – 39) than comparable laboratory studies ^52,56,59^. Specifically, we aimed to recruit a typical sample from previous experiments per input device, so continued recruiting until we had at least n=15 per input device per experimental group.

### Apparatus and Experimental Procedure

Participants completed experiments using their own personal computer (restricted by Prolific’s screening criteria to those using a laptop or desktop) and completed reaching movements in the game using a mouse or trackpad. In the main text we present data from both mouse and trackpad, as we did not observe any meaningful differences between these input devices. Experiments were created using a recently developed framework allowing within and between-participant comparison of Pointing and Looking movements ^15^, which uses the Unity game engine (2019.4.15f) and the Unity Experiment Framework ^72^. The experiments were delivered via a WebGL build hosted on a web page. No calibration procedure was performed to equate stimuli size across disparate participant monitor dimensions, so the physical size of the task was not constant across participants. The experiments were designed to be visible on a 4:3 monitor aspect ratio, with the height always taking up 4 arbitrary Unity units (au) and wider aspect ratios featuring more of a task-irrelevant background texture. The experiments could only be completed in full-screen mode with the desktop cursor locked and hidden, with raw mouse or trackpad input used to perform in-game movements.

Participants first filled in a form to provide details on age, gender, handedness, computer type, and input device (mouse or trackpad), and also clicked a button to ensure game audio was audible. During this stage, participants used their desktop cursor to navigate the form, and the movements of this cursor were tracked in pixel and Unity game units to provide an initial calibration for the in-game cursor. Participants were shown a cut-scene providing exposition for the purpose of the study (popping non-sentient space bubbles), before completing a tutorial that introduced aspects of the experimental task sequentially and interactively to ensure participants fully understood how to complete the experiment. Following this, participants were required to practice using both contexts used in the experiment (Point and Look for Experiments 1-3, or Point and Inverted Look for one group in Experiment 3), where they could switch contexts and adjust cursor sensitivity with key presses. Once participants had completed at least 20 trials in each context, they could progress to the main task.

Across all experiments, participants used their mouse or trackpad to move an FPS style cursor towards a presented target and shoot it (Figure 1a). Throughout a trial, participants saw a dark space themed background. On top of this, the target plane was shown, which had a diameter of 3au, a black background, and a thin ring around it which was white for Experiments 1 and 2 and either yellow or cyan for Experiment 3. When a trial was ready to start, the start-point (0.1au diameter circle centered on the target plane) was colored orange. Participants needed to move their in-game cursor, a small white circle surrounded by a thin white ring of diameter 0.15au (to mimic FPS style reticles), to the start-point and left-click it to start a trial. If participants were within 0.05au of the start-point during this homing movement, the cursor snapped to the start-point to speed up this centering (mimicking FPS-style auto-aim features). Once the start-point had been clicked, a target (0.2au diameter magenta circle) immediately appeared on the target plane, and the start-point turned green to indicate participants could attempt to move to and left-click the target to shoot it. Potential targets were located in increments of 90° on an imaginary circle of diameter 2au. For Experiments 1 and 2, participants had a time limit within which to shoot the target. If the time limit elapsed without a successful shot being registered, the target immediately vanished and a whooshing sound was played. A shot was only successful if, at the time of the left-click, the center of the cursor intercepted the target circle. If the shot was successful, the target exploded and a shooting sound was played. Unsuccessful shots had no effect upon the game, and an unlimited number of attempted shots were allowed per trial. Feedback of a trial outcome, either a successful shot or a lapsed time limit, was provided for 300ms, during which the start-point was colored grey. Upon feedback finishing, the start-point turned orange, indicating that a new trial could begin. Participants had online feedback of the cursor throughout a trial, apart from Experiment 3, where certain trials had no feedback when returning to the start-point.

Cursor movements could be performed in two main contexts, either the Point or Look (Figure 1a, with an Inverted Look context added for Experiment 3, Figure 4a). In the Point context, input movements of the mouse or trackpad translated the cursor while the in-game camera remained static, whereas in the Look and Inverted Look contexts, input movements rotated the camera about a fixed point to change the camera’s view of the scene while the cursor was fixed central to the camera’s view. Critically, the two conditions were equated to ensure identical movements were required to reach a given target in either context. Details of the implementation are described in a previous paper ^15^. Following completion of the experimental trials, participants filled in a questionnaire probing their perception of the experiment’s performance (e.g. lag, difficulty, technical issues), as well as questions about strategy use to overcome the imposed perturbation.

### Experiment 1

Participants (n = 68; mean age = 36, SD age = 11, age range = 20 – 65; 24 male, 42 female) were assigned to complete the experiment in either the Point (n = 32) or Look (n = 36) context. During the main task, participants completed 680 experimental trials. Participants first completed a baseline period of 100 experimental trials where no perturbation was applied to cursor movements. Following this, participants completed 480 experimental trials with a 30° visuomotor rotation, where the cursor movement was rotated away from the actual input movement in either the clockwise or counterclockwise direction (counterbalanced across participants within each group). Finally, participants completed 100 experimental trials where the perturbation was turned off to assess after-effects of learning from the rotation period. Throughout the experiment, trials were tested in cycles of four trials, where each target (0°, 90°, 180° or 270°) was tested in a random order. Participants were given breaks of unrestricted length every 20 cycles through the experiment.

To induce the type of time pressure that might be experienced in a typical FPS-style game, the time limit within which participants had to show a target was continuously staircased throughout the experiment. A pair of staircases, initiated at 450ms and 1050ms for mouse users or 780ms and 1380ms for trackpad users (determined through previous work ^15^), were interleaved and used equally within each block to set the time limit for the upcoming trial. Following a 1-up 1-down procedure, the tested staircase’s time limit was decreased by 30ms following a successful trial, and increased by 30ms following an unsuccessful trial, to approximately give a 50% success rate over the course of the experiment.

### Experiment 2

Participants were assigned to either Experiment 2a (n = 62; mean age = 37, SD age = 11, age range = 18-67; 24 male, 38 female) or Experiment 2b (n = 66; mean age = 36, SD age = 11, age range = 20-75; 20 male, 46 female), and were further assigned to complete the rotation block in either the Point (Experiment 2a, n = 32, Experiment 2b, n = 35) or Look (Experiment 2a, n = 30, Experiment 2b, n = 31) context. The experiment consisted of 800 trials. Participants first completed 100 baseline trials in the ‘untrained’ context (the Look context for the Point Trained group), and then 100 baseline trials in the ‘trained’ context (the Point context for the Point trained group). During these trials, no perturbation was applied to the cursor. Following this, participants completed 480 trials in the trained context with a 30° visuomotor rotation (direction counterbalanced within each group). For Experiment 2a the perturbation was then turned off, and participants completed 100 trials in the untrained context and a further 20 trials in the trained context, to assess the transfer of learning from the trained to untrained context. Participants in Experiment 2b completed the same pair of blocks after the rotation, but the perturbation was left on during these blocks, as a second method of assessing transfer. As in experiment 1, trials were tested in cycles of the four target directions, and the time limit was continuously staircased throughout to induce time pressure. Participants were given untimed breaks after cycles 23 and 45, and thereafter every 20 cycles.

In Experiment 1, it was observed that participants sometimes initiated movements to an uncued target (hereafter ‘jump starts’). To combat this, a further trial outcome was added. If a participant’s movement at a radius of 0.2au was more than 60° in either direction from the ideal movement path between the start-point and target, the trial immediately stopped, and participants were shown an error message for 5 seconds instructing them to make sure they waited to see the target before moving. This check was performed early enough that participants only received a limited sample of their movement on these failed trials. The trial was then repeated, to ensure that participants all received the same number of trials to assess learning and transfer.

### Experiment 3

Participants (n = 76; mean age = 36, SD age = 13, age range = 19-67; 26 male, 49 female) were assigned to use either the Look (n = 40) or Inverted Look (n = 36) context during probe trials. In total, participants completed 760 trials during this experiment. In contrast to the previous experiments, no time limit was used, as pilot testing found this caused a substantial number of errors on the critical probe trials. Further, the ring around the target plane changed color as participants returned their cursor to the start-point to indicate the context of the upcoming trial.

Participants first completed 60 baseline trials to the top and bottom targets (90° and 270°). The first 40 trials had 20 contiguous trials for each tested context (Point and either Look or Inverted Look, in a random order), followed by 20 trials where the two contexts were interleaved to get participants used to switching between contexts. Participants then completed 20 trials in the Point context to the top target, where the no-feedback return was introduced. Here, upon clicking the target, the cursor disappeared, and participants were required to return their hand to a comfortable position within 1.5 seconds of outcome feedback extinguishing. Tones were played after 500ms, 1000ms and 1500ms to ensure participants knew when their hand had to be back to a comfortable position. After the final tone, the cursor appeared in the middle of the start-point. From this trial onwards, Point trials never had return feedback (whereas Look and Inverted Look trials always had return feedback) and were always directed to the 90° (top) target.

Participants then completed the baseline generalization block, an unperturbed version of the generalization block. The baseline generalization block consisted of 15 generalization cycles of 18 trials. The first three trials were in the probed context, either Look or Inverted Look, and probed participants movements to either the left or right target (0° or 180°) in the first trial and both the top and bottom targets in trials two and three (arranged so the top target was tested first either 7 or 8 times, in a random order across cycles). The first movement to a lateral target served as an additional cue, as well as the ring color, that the context had switched before the critical vertical targets were probed. The last 15 trials per cycle were in the Point context to the top target. During the baseline generalization block, movements were unperturbed compared to their nominal mapping. Participants then performed 120 trials in the Point context with a 30° visuomotor rotation applied (rotation direction counterbalanced within each group). Because there was no return feedback, participants only sampled learning during movements to the top target, so should show only local generalization. Participants then completed the generalization block, which had the same structure as the baseline generalization block except the 15 Point trials per cycle had the same visuomotor rotation as in the rotation block (while the probe trials were never perturbed). Because the probe trials had return feedback and probed both trained and untrained directions, unlearning could occur during outward and return reaches, so these 15 trials ‘topped up’ learning to ensure the probes were made after learning again reached asymptotic levels, as determined during pilot testing. While generalization studies typically preclude feedback to avoid this issue, the feedback during Look trials arose because the camera panned and tilted, so clamping the camera’s rotation during Look or Inverted Look probes would give no indication a Look style movement had theoretically occurred. Participants finally completed 20 Point movements with unperturbed feedback, to assess after-effects. Participants were given breaks of unrestricted length during the 8^th^ probe cycle in both the baseline generalization and generalization blocks (after the probe trials but before the top-up trials), and after 100 trials in the rotation block.

The same online check for jump-starts used in Experiments 2a and 2b was used here for trials during the first baseline block for both contexts, and thereafter only for the Point context. As we expected some level of ‘slips’ for the Inverted Look group (i.e. expressions of the non-inverted mapping), we did not check for jump-starts during the probe trials.

### Data Analysis

Kinematic data (cursor position in x and y axis) was sampled at the participant’s screen refresh rate (often approximately 60Hz, 120Hz, or 144Hz) and uploaded to a remote database during the experiments. Offline processing was performed in R (version 4.2.2). Technical issues meant a small number of trials were not uploaded, which removed 0.02% of trials for Experiment 1, 0.05% of trials in Experiment 2, and 0.02% of trials in Experiment 3. Further, a small number of trials were affected by an issue that caused a large single-frame mouse input, causing participant’s cursor to move a large distance suddenly and unexpectedly. This appears to be an issue with WebGL games more broadly, so we could only filter out affected trials post-hoc by removing trials where cursor speed changed by more than 40au/s from a previous or following sample. This removed 0.4% of trials in Experiment 1, 0.04% of trials in Experiment 2, and 0.3% of trials in Experiment 3.

To ensure a common analysis procedure was applied for all participants, movement data was resampled to a consistent 100Hz using linear interpolation and filtered using a second order, zero-phase, low-pass Butterworth filter with a 15Hz cut-off, with the start and end of each trial’s time series padded to avoid transient effects of the filter affecting the movement data. Raw hand paths were visualized (e.g. Figure 2b, rotated to a common target and flipped to show a consistent rotation direction) to ensure the actual movements performed by participants were consistent with canonical observations for these task manipulations. To perform quantitative analysis, key measures were extracted from the time series of cursor movements. The main measure used throughout was hand angle at peak speed. To extract this, the peak radial speed reached during the trial was found and the input position at this time was identified (unaffected by perturbations). Hand angle was then defined as the difference in angle between straight lines connecting the start-point to either the target or input position at peak speed. For participants who experienced a counterclockwise perturbation, the sign of the hand angle was flipped so for both perturbation directions, positive hand angles reflected a movement of the hand in the direction opposite to the perturbation. For Experiment 3, hand angles on Inverted Look trials also had an 180° rotation applied so that they reflected the angle from the ideal aim location to make Look and Inverted Look trials directly comparable. Trials in Experiments 1 and 2 were grouped into cycles of four movements (one to each target), with mean hand angles calculated per cycle, whereas for Experiment 3 hand angles were left per-trial.

In Experiment 1 we noticed participants occasionally initiated movements to an uncued target, likely because the time pressure applied induced some guessing. For example, at extremely low reaction times in forced response paradigms, participants guess uniformly which target will be probed ^73^. To alleviate the effect of this on our data, we removed trials where the absolute hand angle at both peak speed and take-off (hand angle at a radius of 0.2au) were more than 60° from the ideal angle, to ensure the included movements were not directed to an uncued target, which removed 10.5% of trials for the first experiment (*inclusion of these trials did not change any statistical analyses*). To counteract this, Experiments 2 and 3 had online detection of ‘jump starts’ to ensure any such trials were repeated. This led to 5.3% of trials being repeated in Experiment 2, and 1.6% of trials in Experiment 3 (*note the reduction in repeated trials when the time limit was removed for Experiment 3, with a rate comparable to other online adaptation experiments* ^74,75^). Despite this, some trials did not pass the filter at peak speed, removing 1.2% of trials in Experiment 2 (*including these outliers produced identical analyses besides the transfer analysis for Experiment 2b where, despite similar point estimates, there was no statistical difference in transfer between contexts, nor a reduction in learning for the Point-trained group*), and 1.6% of trials in Experiment 3 (*inclusion of these trials did not change any statistical analyses*).

To perform analyses based on periods of the experiment, average hand angles over pre-specified periods were calculated. For Experiment 1, where we wanted to understand how hand angle compared between the contexts across the experiment, the baseline period comprised the last 10 cycles of the baseline block, and early and late rotation and washout were defined as the first and last 10 cycles of the rotation and washout blocks respectively. We also calculated a measure for the after-effect that accounted for potential differences in asymptotic learning by subtracting the hand angle during late learning from that during early washout. For Experiment 2, where we were primarily concerned with transfer, we calculated early and late rotation as in Experiment 1, and additionally calculated transfer as late learning subtracted from the first 10 cycles of the transfer block. For Experiment 3, early and late learning was assessed during the first and last 10 trials during the rotation block respectively. Analysis of probe trials used baseline-corrected values, where the average hand angle per target in the baseline generalization block was subtracted from average hand angle per target in the generalization block for each participant, to correct for any intrinsic biases (analysis was identical with or without baseline correction).

### Statistical Analysis

All statistical analyses were performed in R (version 4.2.2). For all ANOVAs assessing the progression of learning in Experiments 1-3 (performed using the *afex, BayesFactor*, and *bayestestR* packages), in addition to reporting the F-statistic and *p-values*, we included Bayes Factors (BF_10_) as a measure of the evidence for the alternative hypothesis (H_1_) over the null hypothesis (H_0_) to supplement the p-values, and partial eta squared (η_p_2) as a measure of effect size. For all follow-up t-tests using the estimated marginal means (performed using the *emmeans* package) and all independent t-tests, we use two-tailed tests and report the mean and 95% confidence interval of the differences as well as *p-value* and Bayes Factor. Finally, for all mixed-effect linear regressions (featuring a random intercept for participant, performed using the *lme4* package with p-values from the *lmerTest* package calculated according to the Satterthwaite approximation), we report the mean and 95% confidence interval of the regression coefficient. All statistical tests were assessed against a significance threshold of *p* < .05.

## Supplemental videos legends

**Supplemental Video S1:** Video demonstrating experimental trials in Experiment 1 during the Baseline and Rotation phases with Look style visual feedback. Participants attempted to move to and click on a target before a time limit to successfully shoot it, otherwise it disappeared.

**Supplemental Video S2:** Video demonstrating experimental trials in Experiment 1 during the Baseline and Rotation phases with Point style visual feedback. Participants attempted to move to and click on a target before a time limit to successfully shoot it, otherwise it disappeared.

## Data availability

All data and scripts to analyse the data will be made available on the Open Science Framework upon publication of the manuscript.

## Acknowledgements

Authors FM, RM, and MMW are supported by the European Union’s Horizon Research and Innovation programme under grant agreement no. 101070155 and the UKRI through the Horizon Europe Guarantee (#10039307). FM is further supported by a UK Research and Innovation Biotechnology and Biological Sciences Research Council award (BB/X008428/1).

## Author contributions

**Matthew Warburton**: Conceptualization, Methodology, Software, Formal analysis, Investigation, Visualization, Investigation, Writing – Original draft, Writing – Review & Editing. **Carlo Campagnoli**: Methodology, Writing – Review & Editing, Supervision. **Mark Mon-Williams**: Writing – Review & Editing, Funding acquisition. **Faisal Mushtaq**: Writing – Review & Editing, Funding acquisition, Supervision. **J. Ryan Morehead**: Conceptualization, Methodology, Writing - Review & Editing, Supervision, Funding acquisition.

